# Overcoming lentiviral delivery limitations in hard-to-transduce suspension cells for genome-wide CRISPR screening

**DOI:** 10.1101/2025.05.01.651049

**Authors:** Antonino Napoleone, Ivy Rose Sebastian, Federico De Marco, Alexander Molin, Mohamed Hussein, Lovro Kramer, Thomas Jostock, Thomas Kelly, Nicole Borth

## Abstract

Lentiviral vectors are a cornerstone delivery modality of biomedical research, renowned for their ability to stably integrate genetic material into the host genome, enabling sustained transgene expression and long-term genetic manipulation. These properties make them indispensable tools in functional genomics and genome engineering, particularly for delivering molecular components in high-throughput CRISPR screening, a powerful approach for uncovering the genetic basis of complex cellular mechanisms and phenotypes. However, challenges such as lentiviral-induced recombination, unpredictable integration profiles, and variable susceptibility of target cells to transduction can introduce noise and compromise experimental outcomes.

In this study, we selected two suspension-adapted mammalian cell lines, Chinese Hamster Ovary cells CHO-K1 and Human Embryonic Kidney cells HEK293-6E, due to their widespread use in recombinant protein production. Recognizing the influence of intrinsic cell line properties and transduction methodology, we compared two distinct procedures: spinoculation and static transduction. By implementing a two-step static transduction protocol, we achieved significantly higher transduction efficiencies while minimizing cellular stress, streamlining workflows, and eliminating scalability limitations inherent to large-scale lentiviral applications like genome-wide CRISPR screens. To further characterize the variation in lentiviral integration, we used droplet digital PCR (ddPCR) to quantify copy number variation (CNV) both at the pooled population level and within individual clonal isolates.

This comprehensive analysis underscores the robustness of our optimized protocol in enhancing transduction efficiency in difficult-to-transduce suspension cell lines. It further emphasizes the importance of carefully modulating infection rates to limit multiple integrations, ensuring the accuracy and consistency required for large-scale functional genomics applications.

## INTRODUCTION

A central challenge in modern biomedical research lies in developing and refining delivery systems capable of overcoming key limitations in cell and gene therapies, particularly concerning safety, efficiency, and scalability. Both viral and non-viral delivery modalities have been established, each with distinct strengths and limitations^1^. However, their effectiveness relies on application-specific optimization to ensure reliable performance across diverse biological contexts and cell targets. Thus, it is crucial not only to adopt versatile delivery systems but also to tailor and optimize their use to maintain efficacy across various applications. In this scenario, lentiviral vectors have emerged as one of the most robust delivery platforms, offering an unparalleled balance of efficiency and flexibility^2^.

Lentiviruses are retroviruses based on the human immunodeficiency (HIV) type 1 virus and are highly effective at infecting a wide range of dividing and non-dividing target cells^3^. These vectors have been engineered to minimize biosafety concerns by separating essential viral genes across multiple plasmids, ensuring both safety and functionality. Currently, four generations of lentiviral vectors exist, each varying in the number of retained wild-type genes, the inclusion of heterologous elements, and their impact on viral titers and safety. Compared to other viral delivery solutions, lentiviruses have large transgene packaging capacities of up to ∼9 kb^4,5^, and their versatility is particularly enhanced by their pseudotyping capacity, with the vesicular stomatitis virus G (VSV-G) glycoprotein being the most used fusogen for this purpose^6,7^.

Lentiviral vectors are particularly valuable for long-term gene expression studies and genetic manipulation due to their ability to stably integrate genetic material into the host genome^8^. Stable integration ensures sustained transgene expression, making these vectors ideal for applications requiring permanent genetic modifications. Their efficient integration allows for consistent monitoring of gene function, phenotypic outcomes, and the long-term effects of transgene expression, providing a significant advantage over other viral and non-viral delivery systems^9^. For these reasons, lentiviral vectors have become a cornerstone delivery modality in diverse biomedical applications, particularly in functional genomics and genome engineering^10^. A remarkable application is high-throughput genetic screening^11,12^, which harnesses CRISPR (Clustered Regularly Interspaced Short Palindromic Repeats) genome editing technology to interrogate complex cellular mechanisms and uncover the genetic basis of observed phenotypes^13–16^. In recent years, there has been a transformative shift in biological discovery, driven by the effective application of CRISPR screening to uncover unidentified genes, pathways, and regulatory mechanisms^17,18^.

CRISPR screens are typically conducted in a pooled format, wherein thousands of genetically encoded perturbations are simultaneously introduced into cell pools, either on a focused or genome-wide scale^19,20,21^. Individual cells can permanently receive distinct perturbations based on the specific gRNA integrated into their genome^22^. Owing to the stable integrative properties of lentiviral vectors, the abundance of each perturbation can then be quantified by next-generation sequencing (NGS)^23^. In pooled CRISPR screens, tuning the transduction to a low multiplicity of infection (MOI) ensures that most cells receive a single viral integration, as predicted by the Poisson distribution^24^. This setup allows observed phenotypes to be accurately linked to a single gRNA, minimizing the likelihood of multiple perturbations within the same cell, which could otherwise result in ambiguous phenotypic outcomes^25,26^.

Despite these advantages, large-scale genomic screens face several challenges. While streamlined protocols for lentiviral CRISPR screens have been published^23,27,28^, several aspects still require practical consideration and thorough optimization^29,30^. It is important to note that not all cell types respond to lentiviral infection in the same way. Cells differ in their inherent infectivity, and thus transduction efficiency may be notably low in some cell lines known to be harder to transduce, such as primary cells, hematopoietic cells, lymphoid lineage cells, and certain suspension-adapted cells^31–33^. In such scenarios, accurately determining the MOI and the resulting transduction efficiency can be challenging. Variability in the percentage of cells successfully transduced with at least one viral particle can significantly impact the experimental outcome. Furthermore, the lentiviral production process can become extremely tedious and laborious, as larger volumes and high-purity-grade lentiviral preparation is required to attain sufficient transduction efficiency, thus introducing further complexity to the experimental workflow. This poses considerable limitations, especially in the context of genome-wide CRISPR screens.

Likewise, counteracting the low susceptibility to lentiviral infection with tedious and time-consuming methodological adjustments like repeated infection steps, high MOI, or excessive concentrations of transducing units (TU) could potentially lead to heightened cellular stress, diminished viability, or result in multiple transgene copies integrated per cell^34,35^. These conditions have been identified as the most frequent sources of noise in lentiviral-based CRISPR screens^36–38^. Hence, conducting methodical optimization beforehand to pre-empt encountering and troubleshooting these inherent bottlenecks throughout the entire screening process is crucial, mitigating the risk of introducing potential errors that might impact the final output of the screen^39^.

In this study, we focused on two crucial steps in using lentiviral vectors for CRISPR screening: the production phase, during which a library of gRNAs is packaged into the vectors, and the transduction phase, which involves the final delivery into target cells.

We selected two suspension-adapted mammalian cell lines, CHO-K1 and HEK293-6E cells, as model systems for our study because of their industrial relevance and widespread use in recombinant protein production. Chinese Hamster Ovary (CHO) cells are the predominant mammalian system of choice for the manufacturing of therapeutic proteins, mainly due to their capacity to maintain elevated growth kinetics in high-density serum-free suspension cultures, to produce high yields of proteins with human-like glycosylation, and to exhibit a low risk of infection by human pathogenic viruses^40–42^. Conversely, Human Embryonic Kidney293 (HEK) cells have emerged as an attractive and efficient alternative expression host for complex biologics^43–45^ and are considered the gold standard platform for producing viral vectors^46–48^.

Because of their ease of growth and high expression rates, HEK293-6E cells are primarily employed in transient transfection protocols, making them one of the preferred production platforms in the early stages of drug development and pre-clinical studies^49,50^. Both cell lines are currently, and are expected to remain, central to numerous mammalian cell line engineering studies employing CRISPR technology to advance biotherapeutics production and product quality^51–56^.

CHO cells, however, stand out due to their low or absent expression of viral receptor genes, which makes them generally resistant to infection by most viruses^42,57^. The low susceptibility not only minimizes the risk of contamination by human pathogenic viruses, a key reason for their widespread use in industrial biopharmaceutical manufacturing, but it also presents a challenge as it complicates the delivery of transgenes via viral transduction methods.

Therefore, we established an optimized protocol using a third-generation lentiviral system to enhance the transduction efficiency to a level that enables consistent and feasible library delivery, both on a focused and genome-wide scale. These improvements were specifically tailored for a CHO-K1 suspension cell model as a more difficult-to-transduce cell line in comparison to the HEK293-6E cells, used as experimental reference.

Based on the susceptibility to lentiviral infection, largely determined by intrinsic cell line properties and the transduction methodology used, we investigated two distinct procedures, spinoculation and static transduction. Spinoculation is commonly employed to enhance viral infection in difficult-to-transduce cell lines^58–61^. However, it has several inherent drawbacks, including inconsistent results, low scalability, marked cytotoxicity, and time-consuming and laborious workflows. To address these limitations, we established a two-step static transduction protocol as a more effective alternative. Our strategy led to a drastic increase in transduction efficiency, streamlining the experimental timeline, and unlocking any scalability limitation to be expected in genome-wide CRISPR screening.

To validate our approach, we assessed the variation in the integration profile using droplet digital PCR (ddPCR). By simultaneously quantifying the lentiviral vector and the gRNA promoter sequences across the two target cell lines, we were able to monitor the copy number effect and confirm a single integration across the transduced cell population. Importantly, while our protocol was tailored for these specific cell lines, we believe the same adjustments can be successfully applied to other cell lines, whether adherent or in suspension, including those less susceptible to lentiviral infection.

## METHODS

### Cell culture

The HEK293T Lenti-X packaging cell line (Takara/Clontech, cat. 632180) was cultured in Dulbecco’s modified Eagle’s medium (Sigma-Aldrich, cat. D6546), containing 10% fetal bovine serum (Sigma-Aldrich, cat. F7524) and 4 mM L-glutamine (Sigma-Aldrich, cat. G7513), at 37°C and 5% CO_2_ in a humidified atmosphere incubator. Cells were passaged every 2-3 days in T175 tissue culture plates (Greiner Bio-One, cat. 660175) and expanded in Falcon 525cm² 3-layer Multi-Flask (Corning, cat. 11587421) and Falcon 875cm² 5-layer Multi-Flask (Corning, cat. 11597421) for the lentiviral vector production step.

Suspension-adapted CHO-K1 cells (ECACC, 85051005) were cultured in CD-CHO medium (Thermo Fisher Scientific, cat. 10743029) supplemented with 8 mM L-glutamine (Sigma-Aldrich, cat. G7513) and 2 μL/mL Anti Clumping Agent (Thermo Fisher Scientific, cat. 0010057AE) in a humidified atmosphere shaking incubator at 37°C, 7% CO_2_. Suspension-adapted HEK293-6E cells (CNRC, 11565) were cultured in Freestyle F17 medium (Thermo Fisher Scientific, cat. A1383502) supplemented with 4 mM L-glutamine, in a humidified atmosphere shaking incubator at 37°C, 5% CO_2_. Cells in suspension were cultured in TPP Tube-Spin bioreactors (TPP, cat. TPP-87050) and eventually expanded to shake flasks (Corning, cat. 431143) for the scale-up experiments. The cells were passaged every 3-4 days and monitored for viability and viable cell density (VCD) with Vi-CELL™ XR Cell Viability Analyzer (Beckman Coulter). All cell lines used in this study were negative for mycoplasma contamination.

### Vector generation and library amplification

A third-generation lentiviral vector system was employed, involving the co-transfection of four plasmids into packaging HEK293T cells. This included two packaging plasmids: pMDLg/pRRE containing gag and pol, a reverse response element, and the integrase gene, and pRSV-Rev containing an RSV promoter and the *rev* gene, both received from Didier Trono (Addgene plasmids #12251 and #12253, respectively). The pCMV-VSV-G plasmid encoding the vesicular stomatitis virus (VSV)-G glycoprotein gene was purchased from Oxgene (#OG592). The transfer plasmid was derived from the pLL3.7 vector, received from Luk Parijs (Addgene plasmid #11795), into which a CMV-EGFP reporter cassette was introduced for convenient transduction efficiency assessment along with a blasticidin resistance marker (BSD) to enrich for the positively transduced cell population.

The lentiviral guide-expression vector containing two paired guide RNAs (pgRNAs) was constructed in two steps: first, the human U6 promoter and the corresponding gRNA scaffold were cloned in, along with a blasticidin selection marker linked with the EGFP reporter with a T2A linker. Subsequently, the ordered pgRNA oligo pool library (Twist Bioscience) was integrated using a two-step protocol, wherein the pgRNA cassette was first cloned into the modified pLL3.7 base vector via Gibson assembly, followed by the addition of the 7SK promoter and the corresponding gRNA scaffold via Golden Gate assembly. Sanger sequencing was performed to check the integrity of the final plasmid. GeneJET endo-free plasmid maxiprep kit (Thermo Fisher, cat. K0861) was used to purify the plasmid DNA according to the manufacturer’s guidelines. The plasmid DNA concentrations were determined by a Nanodrop Spectrophotometer (Thermo Fisher).

### Lentivirus production

The four lentiviral plasmids, including the pgRNA library vector, the packaging pMDLg/pRRe and pRSV-Rev plasmids, and the CMV-VSV-G envelope plasmid, were co-transfected at a ratio of 1.5:2:1:1, respectively^62^ into HEK293T cells using jetPRIME reagent (Polyplus-transfection, cat. 101000046). The transfection conditions were optimized for the cell line and the cell culture vessels used. A total of 16.5 µg of plasmid DNA amount was transfected in each T175 flask, whereas a total of 49.5 µg and 82.5 µg of DNA amounts were transfected in multi-3-layer and multi-5-layer flasks, respectively. The transfection was performed using a 1:2 DNA to jetPRIME ratio (w/v) according to the instructions provided by the manufacturer. The cells were seeded 24 hours before transfection to get 60-70% cell confluency on the day of transfection. Briefly, the transfection complex was added dropwise onto the cells and distributed evenly. The medium was replaced 4 h after transfection by a fresh cell growth medium supplemented with 0.5 M valproic acid sodium salt (Sigma Aldrich, cat. P4543) to a final concentration of 3.5 mM. Viral supernatants were collected and clarified by centrifugation at 1000 g for 10 min and by filtration with low protein-binding 0.45 µm filter units (Millipore, cat. S2HVU05RE) at 48h and 96h after transfection.

### Lentiviral particles concentration

Viral particles were concentrated by using an in-house 4X lentiviral concentrator solution based on polyethylene glycol (PEG) precipitation^63,64^. To prepare 1000 mL of 4X concentrator solution, 400 g of PEG-8000 (Sigma Aldrich, cat. 89510) and 70 g of NaCl (Carl Roth, cat. 3957.3) were dissolved in 400 mL of MilliQ water and 100 mL of 10X PBS (Sigma Aldrich, cat. D1408-500ML). The mixture was stirred gently until fully dissolved, and the pH was then adjusted to 7.0∼7.2. Subsequently, the buffer was sterile filtered using Stericup® vacuum 0.22 µm filter units (Millipore, cat. SKU X340.1) and stored at 4°C, where it is stable for at least 6 months. Viral concentration is achieved by mixing one volume of 4X concentrator solution with three volumes of clarified lentiviral supernatant. The mixture was collected in 50 mL or 225 mL Falcon tubes (Corning, cat. 352075), gently shaken, and incubated overnight at 4°C. Following that, the mixture was centrifuged at 1600 g for 45 min at 4°C. The supernatant was carefully discarded, and the pellets containing lentiviral particles were thoroughly resuspended in PBS at 1/100 of the original volume. Lentiviral aliquots were immediately stored at -80°C until use. PEG-purified viruses are generally more stable during long-term storage than crude vector preparations.

### Lentiviral titration experiments

#### Spinoculation

For spinoculation, both HEK293-6E and CHO-K1 target cells were seeded in 24 deep-well plates with pyramidal bottom (Enzyscreen, cat. CR1424cl) at a density of 2.5 × 10^6^ cells per well in 0.5 mL of their respective culture medium supplemented with polybrene (Sigma Aldrich, cat. TR-1003-G) at a final concentration of 8 µg/mL. To assess the effects of varying lentiviral concentrations on each cell line, two different gradient dilution series were prepared, along with non-infected controls (NIC) that received no virus. Concentrated lentiviral aliquots of 250 µL, 125 µL, 85 µL, 50 µL, and 25 µL were diluted in 0.5 mL of the respective culture media for CHO, and aliquots of 250 µL, 125 µL, 50 µL, 25 µL, and 12.5 µL for HEK293-6E. No anti-clumping agent was used during the transduction process. The cells were then incubated with the concentrated lentiviral preparation for 15 minutes at room temperature. Following this, plates were centrifuged at 800 g for 2 hours at 32°C. Subsequently, 1.5 mL of respective cell culture media without polybrene was added dropwise to each well to a final volume of 2.5 mL. The cells were gently resuspended before being incubated overnight at 37°C. The following day, the cells were spun down, resuspended in fresh growth medium, and transferred into TubeSpin® Bioreactor 50 mL tubes (TPP, cat. 87050), back to shaking conditions.

#### Two-step static transduction

For the *two-step static transduction*, HEK293-6E and CHO-K1 target cells were seeded in 6-well plates with flat bottom (Greiner Bio-One, cat. GRE-657160) at a density of 2.5 × 10^6^ cells per well in 0.5 mL of their respective culture media supplemented with polybrene at a final concentration of 8 µg/mL. As above, to assess the effects of varying lentiviral concentrations on each cell line, two different gradient dilution series were prepared as described above, along with NIC. No anti-clumping agent was used during the transduction process. The cells were incubated with the concentrated lentiviral preparation for 15 minutes at room temperature and then transferred to a static incubator at 37°C for 16 hours (overnight incubation). Following that, 1.5 mL of respective cell culture media without polybrene was added dropwise to each well to a final volume of 2.5 mL, to mitigate the cytotoxic and antiproliferative effect of polybrene. The cells were gently resuspended before being transferred back to a static incubator at 37°C. At 24 hours post-transduction, the cells were spun down, resuspended in fresh complete medium, and transferred into TubeSpin® Bioreactor 50 mL tubes in the shaking incubator.

### Functional titer determination and MOI calculation

The functional titers are defined by the number of infectious virions required to transduce an individual cell in a defined volume. Functional titers were determined by evaluating the transduction efficiency through the quantification of EGFP-expressing cells using flow cytometry at 4 days post-transduction. Flow cytometry was performed using a CytoFLEX (Beckman Coulter, USA), and the obtained data were analyzed using Kaluza Analysis Software version 2.1. The non-infected controls (NIC) were used as negative controls to set the gate for the EGFP-positive cells during the analysis. This approach aids in identifying the ideal experimental conditions to reach the desired efficiency range and ascertain the optimal MOI tailored to each target cell line.

The lentiviral functional titers or transduction units per mL (TU/mL) were calculated as follows:

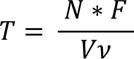

Where:

○ T = Infectious Titers, TU/mL
○ N = Number of cells transduced
○ F = Percentage of cells expressing fluorescent marker (GFP)
○ Vν = Virus volume, mL

The MOI was calculated as an endpoint measurement following the quantification of the transduction efficiency in each target cell line, and applying the Poisson distribution as follows:

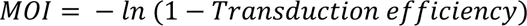

### CRISPR screen on a targeted scale

CRISPR screens were conducted exclusively using the *two-step static transduction* protocol, employing a pilot library containing 2000 pgRNAs. HEK293-6E and CHO-K1 cells were seeded in 6-well plates with flat bottoms at a density of 2.5 × 10^6^ cells per well in their respective culture media supplemented with polybrene at a final concentration of 8 µg/mL. Based on the previous titration experiments, the transduction conditions were tailored to achieve an efficiency range of 14-16% (MOI ≈ 0.16) in both target cell lines, minimizing the likelihood of multiple integrations. A higher volume of lentiviral vector was required for effective transduction in CHO cells compared to HEK293 cells, with 25 µl per well used for CHO cells and 12.5 µl per well for HEK293 cells. Consistency in library delivery was ensured by adhering to the experimental parameters established during lentiviral titration experiments. Transduction of 15 × 10^6^ cells was achieved by pooling six wells of a 6-well plate per replicate following the *two-step static transduction* procedure, after which cells were transferred into 125 mL flasks (Corning, cat. 431143) and returned to shaking conditions. Transduction efficiency was assessed after 4 days by flow cytometric analysis of eGFP-expressing cells compared to NICs. Details regarding the calculation of the number of cells required to maintain library representation at the time of lentiviral delivery are provided in the accompanying table following the formula:

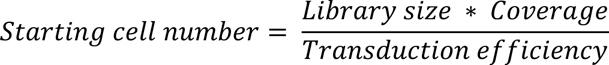

| Calculation of the number of cells needed to maintain library representation |  |  |  |
| --- | --- | --- | --- |
| Library size | Coverage | Transduction efficiency | Starting cell number |
| 2000 pgRNAs | 1200X | 16% | **15,000,000** |

### Genome-wide CRISPR screen

A genome-wide CRISPR screen was conducted to validate the scalability of our two-step static transduction protocol, delivering a library containing 112,000 pgRNAs. The lentiviral library was generated using the Lenti-X™ HEK293T packaging cell line. Eight multi-5-layer flasks were used to produce the genome-wide lentiviral library, employing the jetPRIME transfection protocol as described above. Viral supernatants were harvested at 48- and 96-hours post-transfection, then clarified by centrifugation and

0.45 µm filtration. Subsequently, the viral supernatant underwent concentration using the 4X lentiviral concentrator solution via PEG precipitation, resulting in stocks with 100-fold concentration. Three independent transductions were executed in CHO-K1 cells, which showed a lower susceptibility to lentiviral infection compared to HEK293-6E cells. To accommodate a larger number of transduced cells per condition, the two-step static transduction protocol was upscaled from a standard 6-well plate format to a larger 145 mm cell culture dish (Greiner-Bio-One, cat. 639160). This upscaling involved adjusting various parameters such as transduction medium volume, initial cell number, and lentiviral dilution in proportion to the 15-fold increase in the cell dish surface area. The transduction conditions were tailored to achieve a 14-16% transduction efficiency range (MOI ≈ 0.16), to limit the likelihood of multiple lentiviral integrations. A total of 37.5 million cells were seeded into each 145 mm cell culture dish in 7.5 mL transduction medium containing polybrene at a final concentration of 8 µg/mL. One aliquot of concentrated lentiviral stock was prepared in 7.5 mL and added to the cells, based on the lentiviral titration experiment previously determined in CHO-K1 cells. Following that, fresh cell culture media without polybrene was gently added to each cell dish to a final volume of 25 mL. At 24 hours post-transduction, cells were resuspended in fresh media and transferred to shaking conditions in 1 L flasks (Corning, cat. 431147). Transduction of 375 MVC was achieved by transducing ten cell culture dishes per screen sample to retain 500-fold library coverage. Transduction efficiency was assessed after 4 days by flow cytometric analysis of GFP-expressing cells compared to NICs.

| Calculation of the number of cells needed to maintain library representation |  |  |  |
| --- | --- | --- | --- |
| Library size | Coverage | Transduction efficiency | Starting cell number |
| 112,000 gRNAs | 500X | 16% | **350,000,000** |

### Genomic DNA isolation

Following the lentiviral transduction, the target cells were selected with 10 μg/ml blasticidin for 10 days until full recovery. Genomic DNA (gDNA) was extracted from both target cell lines, using the DNAeasy Blood & Tissue kit (Qiagen, cat. 69506) according to the manufacturer’s guidelines. The isolated DNA concentrations were determined by a Nanodrop Spectrophotometer (Thermo Fisher Scientific).

### Quantification of copy number using ddPCR

*Primer design:* Primer pairs were designed after checking for target specificity in both the human and CHO genomes (PrimerBLAST, NCBI) and were ordered from IDT (Integrated DNA Technologies) (Table 1).

**Table 1.**
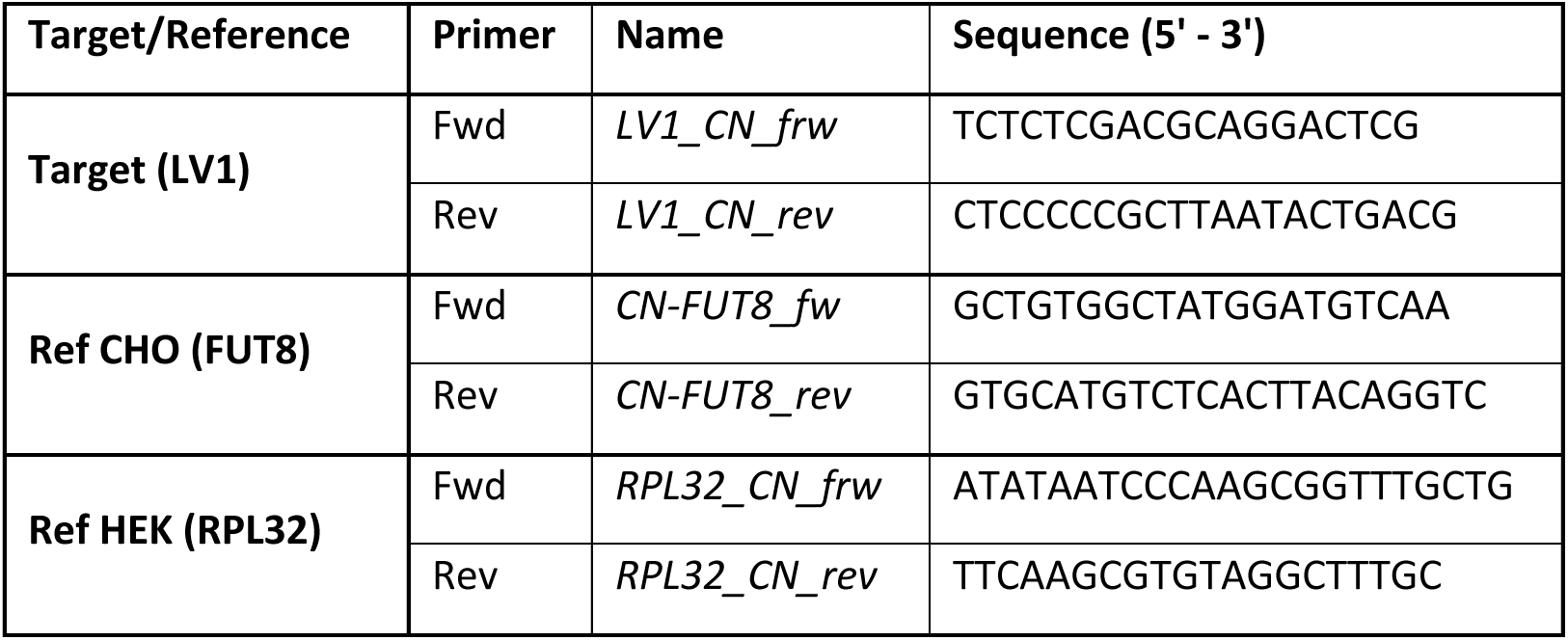
Primer sequences used for copy number quantification by ddPCR.

*ddPCR run and data analysis:* For the copy number determination, the QX200 Droplet Digital PCR System (Bio-Rad Laboratories, Inc.) was used in combination with the EvaGreen DNA-binding dye (Bio-Rad, cat ID, 1864033) according to the manufacturer’s guidelines unless stated otherwise.

Prior to the ddPCR run, 1 µg of the gDNA samples was digested using HindIII in a final volume of 50 µL, after verifying that the enzyme did not cut within the amplification area. The final ddPCR master mix was prepared using 1X QX200™ ddPCR™ EvaGreen Supermix, reverse and forward primers of the reference or the target sequences, respectively, at final concentrations of 0.25 µM, an input of 100 ng of the respective pre-digested gDNA, and nuclease-free water in a total volume of 22 µL. The ratio of master mix to input DNA was kept at 1:4. These mixes were then loaded into the DG8™ Cartridge (Bio-Rad, cat ID 1864008), and droplets were generated using the QX200™ Droplet Generator (Bio-Rad Laboratories, Inc.). The generated droplets were then carefully loaded into a clean PCR plate, heat-sealed for 5 s at 180°C using a PX1 PCR plate sealer (Bio-Rad Laboratories, Inc.), and loaded into the C1000 Touch thermal cycler with a 96–deep well reaction module (Bio-Rad Laboratories, Inc.). The amplification was performed with a ramp rate of 2°C/sec and the following cycling conditions: initial denaturation at 95°C for 5 min, 40 cycles at 95°C for 30 s and 60°C for 1 min, followed by a signal stabilization step at 4°C for 5 min, and a final step at 90°C for 5 min. Data acquisition and analysis were performed with the QX200™ Droplet Reader and QuantaSoft™ Software (Bio-Rad Laboratories, Inc.). The target copy number for all conditions was assessed by running PCRs for both target and reference genes (FUT8 and RPL32 for CHO and HEK, respectively) for each of the transduction conditions. The copy number was calculated from the absolute ddPCR quantification value obtained as concentration (copies /µL), using the following formula:

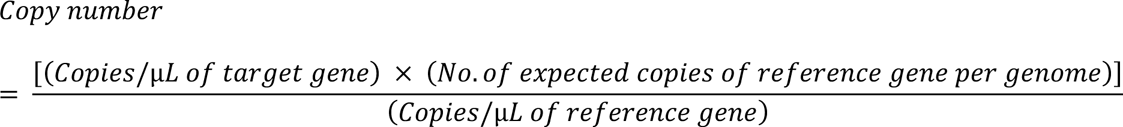

For FUT8 and RPL32, the number of expected copies per genome was two.

### Statistical analysis

All measurements were done as independent replicates. Sample means ± SD of four independent replicates were calculated for the lentiviral titration experiment. To determine the significance of differences between two groups, the one-sided Mann-Whitney U-test for independent samples was applied. Statistical analyses were conducted using R software (v4.3.1). A p-value of <0.05 was defined as being significant: *P < 0.05; **P < 0.01; ***P < 0.001; ****P<0.0001.

## RESULTS

To ensure consistent and reproducible library transduction, we focused on four key experimental phases:

1) Optimizing the lentiviral four-plasmid co-transfection protocol, scaling it up in multi-layer flasks.
2) Increasing viral titers through a cost-effective viral concentration step prior to transduction.
3) Refining the lentiviral transduction methodology to enhance efficiency and consistency and reduce cellular stress in hard-to-transduce suspension cell lines.
4) Streamlining the genome-wide screen upscaling conditions without compromising cell viability or overcomplicating the experimental workflow.

The enhanced performance of our optimized transduction protocol for efficient CRISPR library delivery was validated on both a small, targeted scale and a genome-wide scale.

### 1. Upscaling of high-yield lentiviral packaging in multilayer flasks with valproic acid enhancement

The lentiviral packaging step was optimized by fine-tuning and upscaling the transient transfection conditions from standard T flasks to 3- and 5-layer cell culture flasks. A key challenge in large-scale lentiviral production is maintaining consistent functional titers while reducing batch variability and minimizing reagent costs. This was achieved by transfecting in 3-layer and 5-layer flasks, respectively, which yielded lentiviral titers comparable to those achieved in individual T flasks. This upscaling strategy not only maintained the quality and functionality of the lentiviral particles but also significantly reduced the number of individual transfections required, thereby minimizing batch-to-batch variability.

Using this transfection protocol, similar or improved yields were achieved while using lower concentrations of plasmid DNA and jetPRIME reagent. Furthermore, the use of valproic acid (VPA) at a final concentration of 3.5 mM^65,66^ prolonged the transgene expression by reducing the replication rate of the packaging cells and stabilizing the cell culture environment following transfection. This enabled the harvesting of two lentiviral batches at 48 and 96 h post-transfection, both yielding comparable quantities of functional viral preparations (data not shown).

### 2. Efficient and cost-effective lentiviral concentration using PEG precipitation

To obtain higher infectious titers required for difficult-to-transduce cell lines, we integrated a lentiviral concentration step into our pipeline using PEG precipitation. PEG precipitation is an efficient and inexpensive method, with a straightforward and time-effective protocol that does not require specialized equipment, making it easily accessible in BSL-2 lab settings. This procedure allowed the removal of contaminants such as cell debris and exhausted serum-containing medium without any need for ultracentrifugation steps, which can sometimes negatively impact viral titers^33^. The main advantage in the context of genome-wide screens is its easy scalability to large volumes, leading to high recovery rates of infectious viruses. Concentrated lentiviral particles obtained using PEG-8000 precipitation are typically stable and can be stored for extended periods with minimal loss in infectious titers. This implementation allowed us to achieve up to 100-fold concentrated viral stock relative to the raw supernatant. Additionally, this approach enhanced overall cost-effectiveness, with the estimated cost of preparing the (4X) lentiviral concentration solution in-house being approximately 10- to 20-fold lower than that of commercially available options.

### 3. Refining the transduction procedure for enhanced lentiviral delivery

Transduction dynamics vary among different cell types, primarily influenced by intrinsic properties specific to each cell line, which may dictate susceptibility to lentiviral infection. Moreover, the kinetics of infection slightly vary according to the experimental conditions and the transduction medium used. The presence of polybrene can play a crucial role in influencing these kinetics, as it is often employed to enhance infection efficiency by overcoming the repulsive electrostatic forces between the cell membrane and the viruses^67^. Nevertheless, retroviruses, including lentiviruses, can only travel a limited distance through random Brownian motion in the infection medium until they adsorb onto target cells. The probability of adsorption is expected to be inversely proportional to the distance between the virus and the target cells, while the time required for capture is proportional to the square of that distance^68^.

This probability is further affected by the fact that suspension cells are maintained in a three-dimensional environment, where contact with the virus is limited by the increased surface-to-volume ratio compared to 2-D cell cultures^69^. Therefore, the first parameter to consider was minimizing the physical distance between the viruses and the target cells in suspension. This involved concentrating viral stocks using PEG-based precipitation to reduce the transduction media volume and increasing target cell density to optimize the number of cells for transduction, based on the experimental scale. High cell density has been shown to enhance transduction efficiency with lentivirus^70^. The two methods used were spinoculation and an optimized two-step static transduction protocol.

#### Spinoculation

Spinoculation is a commonly used method to enhance the infection rate in suspension-adapted and other hard-to-transduce cell lines. Different studies have tried to elucidate this mechanism, where some hypothesize that low-speed centrifugation may generate a concentrating effect that aids viral penetration, although the gravitational force is too low to allow for virus sedimentation^59^. Others have suggested that centrifugation may increase the binding capacity of the virus to cell membranes in a liquid solution or exert some biochemical effects on cells, rendering them more susceptible to infection^61^. However, these mechanisms remain to be elucidated.

We noticed that, under identical experimental conditions, the transduction efficacy was not always comparable or reproducible. This methodology proved to be stressful for the cells, affecting the overall cell viability and slowing down cell recovery post-transduction for several days. These findings align with the observations reported by Rajabzadeh et al., particularly when spinoculation is used in conjunction with polybrene, resulting in decreased consistency in transduction efficiency within the cell population. Furthermore, spinoculation presents a highly laborious and time-intensive procedure that adds complexity and delays the screening workflow. The limited scalability, the necessity for skilled personnel, adequate equipment, and cell culture vessels, combined with the lengthy pipetting procedures, can render this methodology a challenge.

#### Two-step static transduction

Hence, we established and refined a method known as two-step static transduction to enhance the transduction efficiency while streamlining the experimental timeline and reducing the complexity, irrespective of the cell type or the number of cells to transduce. The static transduction experiments were carried out in two-dimensional culture vessels, with the infection medium added to the target cell bed. In a 6-well plate, the target cells were initially exposed to 1 mL of transduction medium containing the concentrated viral preparation and polybrene for 16 hours (overnight incubation). Following the first step, 1.5 mL of standard growth medium (without virus and polybrene) was added to a final volume of 2.5 mL to mitigate the detrimental cytotoxic and antiproliferative effects of polybrene^71^ and accelerate cell recovery. Maximum transduction efficiency was observed within 34 hours, aligning with the estimated half-life of VSV-G enveloped lentiviral vectors, as reported by Dautzenberg *et al.*^6^. Consequently, we extended the transduction period to 24 hours, after which no substantial further increase in efficiency was observed. The lentiviral solution was then fully replaced with standard growth medium, and the cells were returned to shaking conditions. These methodological adjustments not only maximize efficiency but also streamline the transduction procedure, reducing the need for extensive manual intervention and mitigating cell stress. The overall workflow of the optimized protocol is presented in Figure 1.

**Figure 1.**
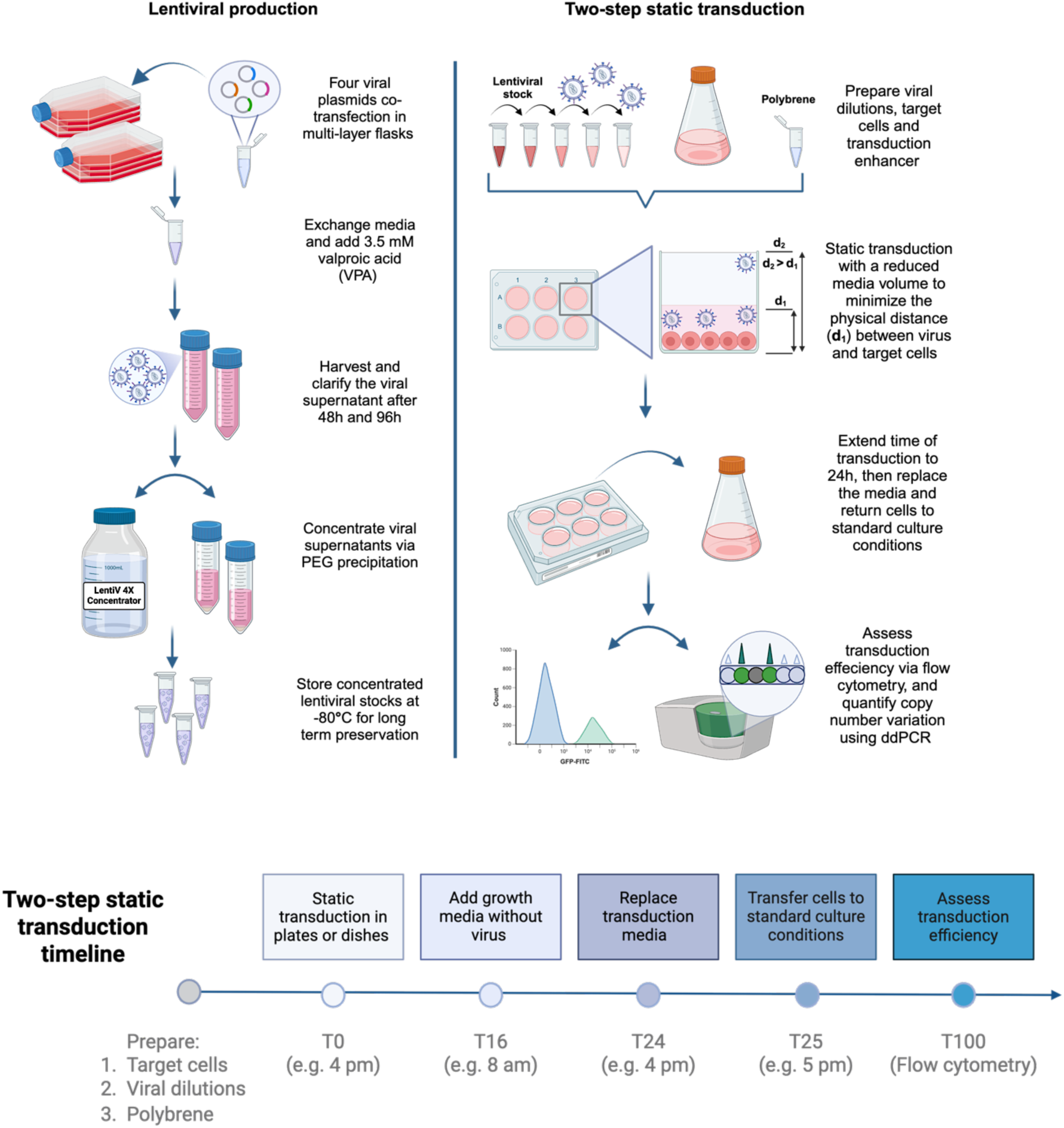
Schematic overview and experimental timeline of the lentiviral production and the two-step static transduction protocol. The optimized production pipeline enables efficient and scalable generation of high-titer lentiviral stocks. During transduction, the distance between the viral particles and the target cell bed influences the viral adsorption probability. Retroviruses, like lentivirus, travel an average distance within one half-life in infection medium by Brownian motion over the transduction period, as demonstrated by Chuck et al. Decreasing the transduction volume and extending the transduction time to 24 hours significantly enhanced the transduction efficiency, streamlining the procedure and facilitating scalability. Created in BioRender. N, A. (2025) https://BioRender.com/42cxuwc.

#### Improved efficiency in lentiviral transduction with two-step static transduction

The primary step toward effective lentiviral library delivery involves determining the infectious titers. Conventional assays based on the p24 lentiviral capsid antigen detection are intended to measure the non-functional or physical titers. These assays usually tend to give overestimated quantifications due to the detection of free p24 antigen in the viral supernatant or the presence of defective virions produced during the packaging step. To address these limitations, quantification of the functional titers was performed through reporter gene expression analysis or transduction efficiency by flow cytometry on the transduced cell population subjected to gradients of lentiviral dilutions.

The target cells were transduced using a 100-fold concentrated lentiviral preparation, applied in five gradient dilutions. To ensure consistency, the same initial cell number was used for all transductions, with cells seeded at a density of 2.5 × 10⁶ per well. Spinoculation was performed in 24-deep well plates with a pyramidal bottom, while static transductions were carried out in 6-well plates.

A significant enhancement in transduction efficiency was observed with the optimized two-step static protocol (Figure 2). This approach facilitated the transduction of an equivalent initial cell number, effectively diminishing the overall experimental duration and speeding up cell recovery after transduction while achieving nearly double the transduction efficiency compared to spinoculation. These findings remained consistent across all gradient dilutions examined and in both target cell lines. As expected, we observed that CHO-K1 cells exhibited lower susceptibility to lentiviral infection compared to HEK293-6E cells.

**Figure 2.**
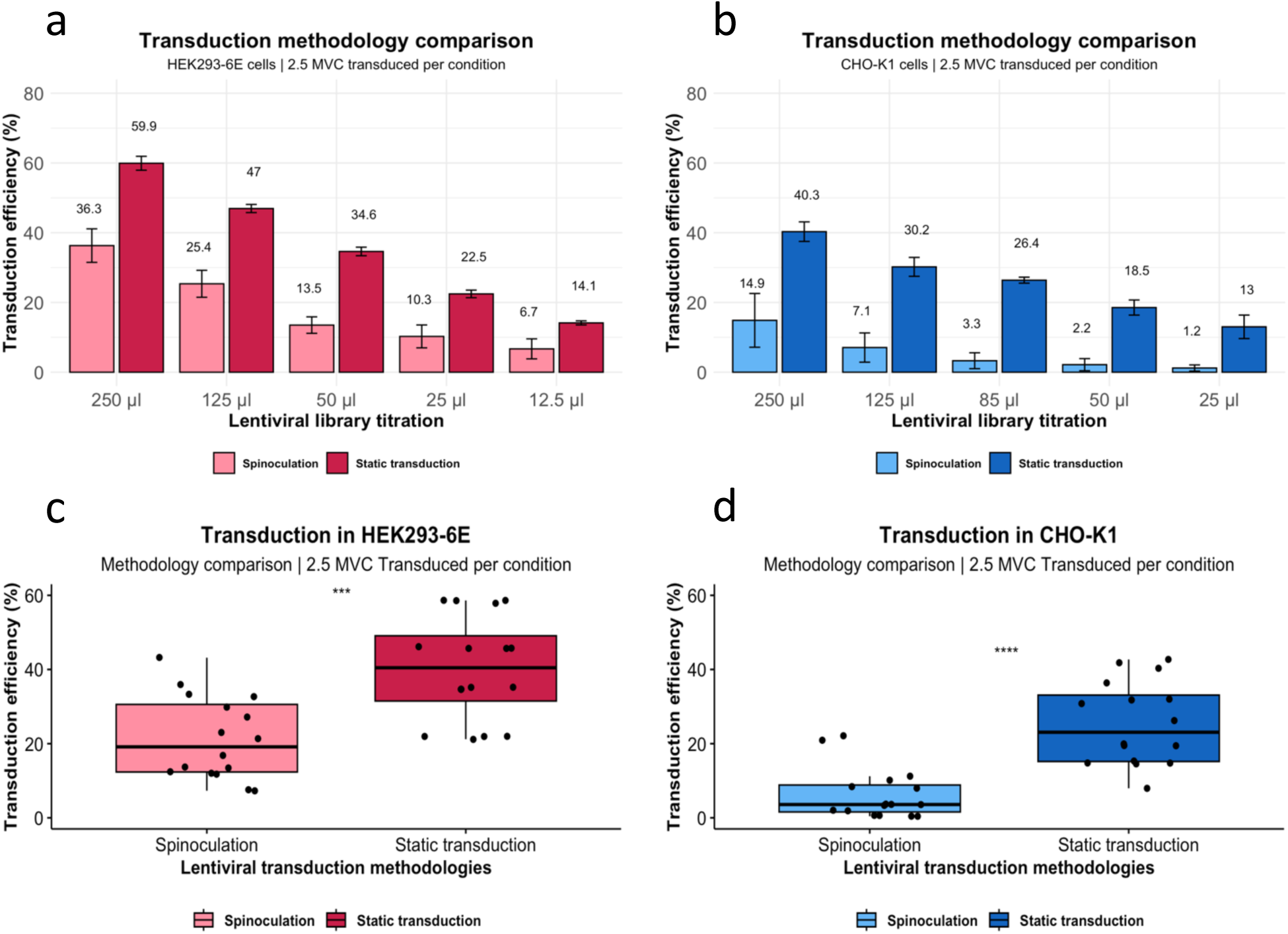
Comparison of transduction methods: static transduction versus spinoculation. Transduction efficiency, measured as the percentage of cells expressing eGFP, was quantified by flow cytometry. Each condition was tested in four biological replicates, and results were normalized to non-infected controls (NIC). Efficiency analysis of spinoculation was performed on HEK293-6E cells and CHO-K1 cells across five distinct conditions with gradient dilutions of the lentiviral library (Figures 2a and 2b). The two-step static transduction protocol demonstrated significantly higher efficiency compared to spinoculation across both HEK293-6E and CHO-K1 cell lines (Figures 2c and 2d). Error bars in Figures 2a and 2b represent the standard deviation of four biological replicates. The data shown in Figures 2c and 2d represent grouped efficiency results from the transduction experiments. Statistical significance was defined as *P < 0.05; **P < 0.01; ***P < 0.001; ****P < 0.0001.

### 4. Achieving consistent and accurate lentiviral transductions in a targeted CRISPR screen

After confirming improved transduction efficiency, we validated lentiviral library delivery by performing a pooled CRISPR screen at a targeted scale in both cell lines. Consistent library delivery was ensured by the experimental parameters established during lentiviral titration, considering that CHO cells required twice the viral particle concentration for effective transduction. Using the optimized two-step static transduction methodology, we delivered a pilot library of 2000 guides, achieving the desired transduction rate of 14-16% (MOI ≈ 0.16). This balance between efficiency and simplicity ensured adequate library representation, avoiding excessive increases in cell numbers or transgene copy number integrations. The adjustment allowed us to maintain a 1000-fold library representation while achieving the target transduction efficiency range under sixfold scaled-up conditions. The methodology demonstrated high reproducibility and consistency, ensuring controlled transgene delivery with minimal variation across independent transduction experiments (Figure 3). Droplet digital PCR (ddPCR) analysis confirmed a predominantly single-copy integration in both cell lines, with an average copy number of 1.09 in CHO-K1 and 1.06 in HEK293-6E, further validating the accuracy and robustness of the delivery system for downstream functional screening. Furthermore, this approach shortened the experimental timeline by eliminating labor-intensive steps and reducing cellular stress compared to spinoculation. Overall, this strategy enables efficient and scalable lentiviral transduction for pooled CRISPR screens, offering a reliable and streamlined approach for large-scale experiments.

**Figure 3.**
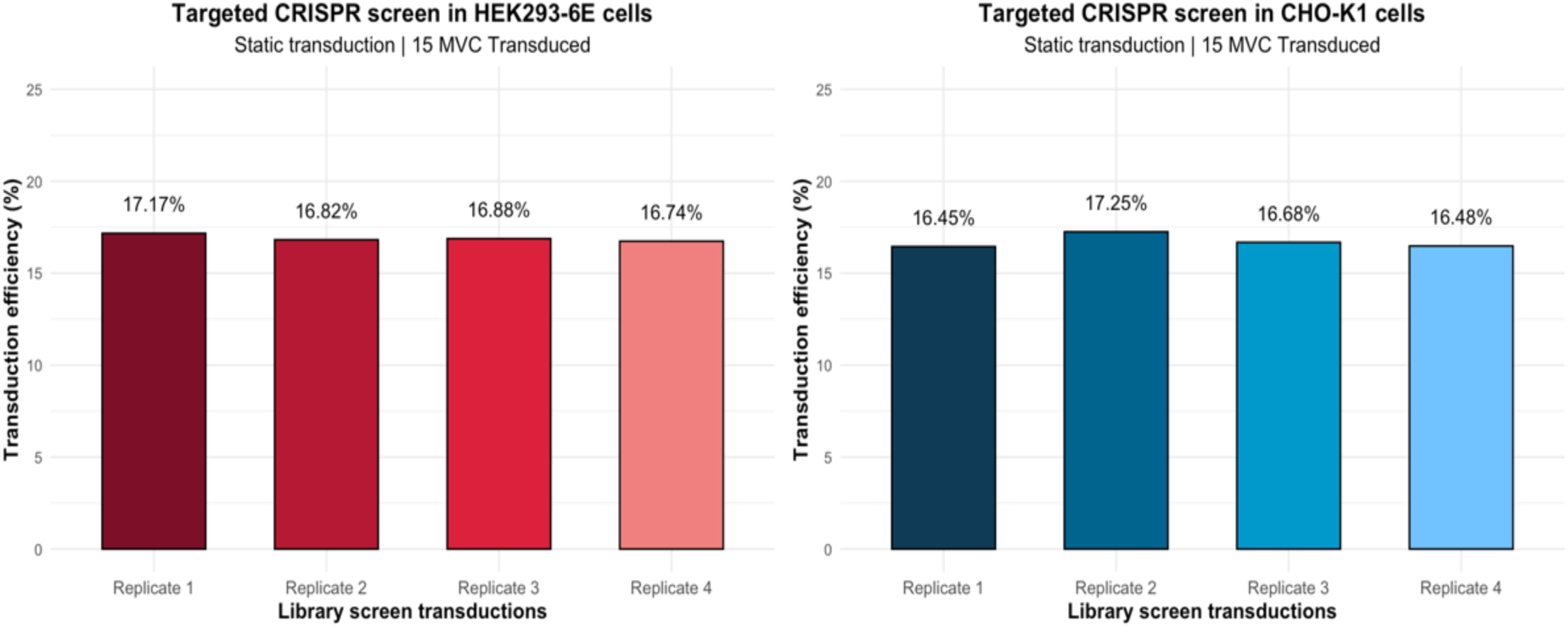
Targeted CRISPR screen transductions. Four independent transductions were performed in CHO-K1 and HEK293-6E suspension cell lines using the two-step static transduction methodology. **Left**: CHO-K1 cells and **right**: HEK293-6E cells were transduced with a target transduction rate of 14-16% (MOI ≈ 0.16), and transduction efficiency was quantified as the percentage of eGFP-expressing cells via flow cytometry. Efficiency values were normalized to non-infected controls (NIC).

### 5. Maximizing genome-wide CRISPR screen scalability with controlled copy number variation

Next, a custom genome-wide library was designed comprising 112,000 pgRNAs, targeting specific genomic regions within the CHO-K1 cell genome. Lentiviral production was upscaled into 5-layer flasks to meet the demands of a large-scale library delivery. To achieve a controlled transduction efficiency of 14-16% (MOI ≈ 0.16) with 500-fold coverage, at least 3.75 × 10^8^ cells needed to be transduced. To accommodate this scale, the experiment was conducted in ten parallel 145 mm cell culture dishes instead of 6-well plates, enabling synchronous transfection of 3.75 × 10^7^ cells per dish. The transition was guided by the surface area ratio, requiring a 15-fold increase in cell numbers, transduction media, and lentiviral volumes. Scaling up to 145 mm dishes enabled transduction of a sufficient number of cells in ten dishes, performed in three independent experiments, maintaining transduction efficiency, workflow simplicity, and without inducing cellular stress (Figure 4a). To ensure that the majority of the transduced population contained single-copy integrations, a ddPCR verification step was included to quantify the copy number variation (CNV) both across the entire cell pool and within individual clonal isolates and to assess how closely the results aligned with a Poisson distribution. By analyzing the transduction outcomes at several lentiviral titrations, we observed a correlation between transduction efficiency and average copy number. At lower transduction efficiencies (14-16%, MOI ≈ 0.16), the average copy number was 1.4, indicating single-copy lentiviral integrations within the transduced cell population (Figure 4b). This MOI reflects the optimal range for pooled CRISPR screens, where minimizing multiple integrations is critical for maintaining data interpretability. Conversely, higher transduction efficiencies (40-42%, MOI ≈ 0.54) were associated with elevated copy numbers (1.9) and a higher likelihood of multiple perturbations.

**Figure 4.**
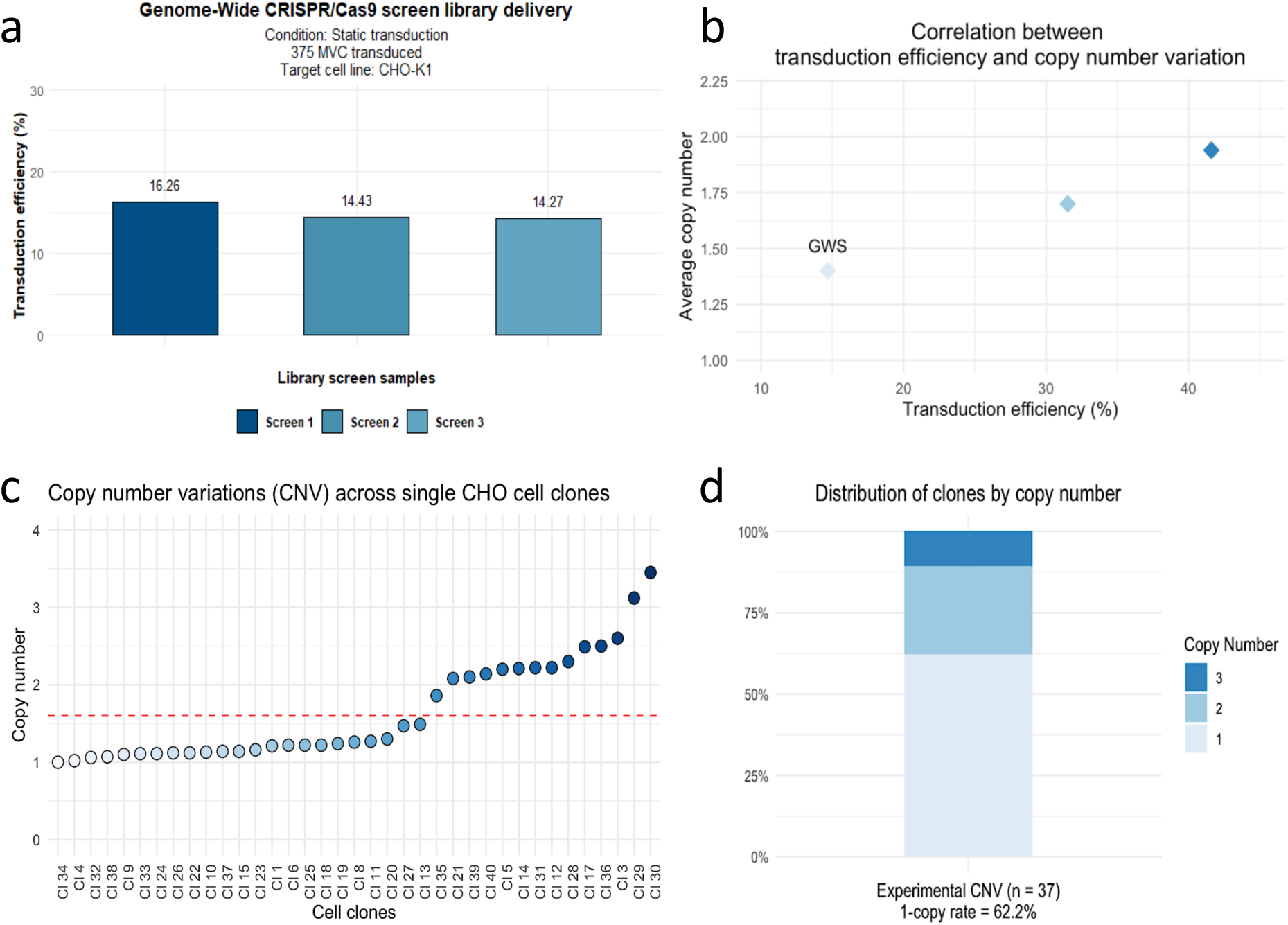
Genome-wide CRISPR screen transduction experiments and copy number analysis in CHO-K1 cells. **(4a)** Three independent experiments were conducted while scaling up the two-step static transduction protocol into 145 mm cell dishes to achieve the required number of transduced cells. With a target value of 14-16% transduction efficiency (MOI ≈ 0.16), this approach enabled the transduction of 3,75 × 10^8^ cells per condition, ensuring a library representation of 500-folds at the time of transduction. **(4b)** ddPCR analysis of a lentiviral library titration experiment reveals a correlation between increasing transduction efficiency and higher average copy number. The labeled data point (“GWS”) highlights the condition selected for the genome-wide screen, corresponding to an average copy number of 1.4 to ensure predominantly single-copy integrations. **(4c)** Copy number distribution in individual subclones (n = 37), analyzed 21 days after single-cell sorting of the transduced population, reveals a skewed pattern favoring single-copy integration. The dashed red line at y = 1.6 represents the average copy number across all individual cell clones, correlating closely with the average copy number measured at the cell pooled population level (1.4). **(4d)** The bar plot illustrates the percentage distribution of single-cell clones with different copy numbers (1, 2, and 3).

To further evaluate the vector copy distribution in the transduced population, single cells were isolated using fluorescent-activated cell sorting (FACS), based on the expression of the eGFP reporter encoded in the plasmid library vector. Genomic DNA from 37 expanded clones was analyzed by ddPCR, revealing a skewed copy number distribution towards single-copy integration. Among the subclones, 23 contained a single copy (62.2%), 10 had two copies (27%), and 4 had three or more copies (10.8%), closely correlating with the average copy number observed in the pooled population (Figure 4c-d).

## Discussion

Forward genetic screens using the CRISPR toolbox have emerged as a powerful strategy for unraveling gene function and intricate molecular mechanisms within mammalian cell genomes^25,72^. This approach has greatly accelerated the identification of pivotal genetic targets for a variety of genome editing applications^73,74^, making it indispensable for advancing mammalian cell line engineering^75–77^. Such engineering approaches hold considerable promise for further enhancing growth kinetics, recombinant protein expression, and quality, rendering them a highly sought-after field within biopharmaceutical research^16,40,54,78^.

Although CHO cells are highly favored in industrial biopharmaceutical manufacturing due to their high productivity, ability to grow at high densities in suspension, and strong regulatory acceptance^79^, their low expression of viral receptor genes makes them resistant to most viral infections^42,57^. This resistance, while advantageous for ensuring the safety of biotherapeutics, poses unique limitations for effective viral vector-based transgene delivery. Similarly, lentiviral vectors, which are the preferred system for packaging and delivering CRISPR screen components^80^, face significant challenges when transducing certain cell types, such as suspension lines, particularly when attempting to achieve consistent and efficient results at scale^22^. Suspension cell lines have a smaller surface area compared to adherent cells, which makes it harder for lentiviral particles to interact with the cell membranes and efficiently enter the host cells. In addition, suspension cells grow in three-dimensional cultures, reducing the probability for virions to come into contact with the target cells compared to adherent cell lines.

Our study focused on optimizing two central aspects of the pooled CRISPR screen workflow: the lentiviral library packaging and the transduction procedure. These implementations were particularly tailored to overcome the challenges associated with the transduction of CHO-K1 suspension cells, a model that we compared with the more transduction-permissive HEK293-6E cell line, used as an experimental reference.

To improve lentiviral production, we optimized the transfection process by scaling up the jetPRIME protocol into multilayer flasks. This bulk approach minimized the variability inherent in multiple individual transfections, improving batch-to-batch consistency while reducing the overall usage of plasmid DNA and jetPRIME reagent. Importantly, these adjustments did not compromise viral yields, making the process more cost-effective and efficient. The addition of valproic acid (VPA), consistent with previous reports demonstrating that histone deacetylase inhibitors improve transfection protocols^62,65,66,81^, extended the lifespan of transfected cells so as to enable two consecutive harvests. Together, these optimizations led to consistent and homogenous yields across all viral production batches, increasing the final titer to address the critical need for reproducibility and scalability in high-throughput applications. Finally, incorporating PEG8000-based precipitation for viral concentration streamlined the workflow and increased infectious titers without the need for ultracentrifugation. This yielded better transduction outcomes and ensured the stability of our concentrated viral stocks during prolonged storage, even after 15 months.

To enhance the transduction methodology for the difficult-to-transduce CHO-K1 cells, two protocols were compared. While spinoculation has been extensively used to enhance the viral infection rates in difficult-to-transduce cell lines^58–60^, it poses significant challenges in terms of reproducibility, cell recovery, and scalability^61^. Our evaluation of transduction methodologies highlighted the superiority of the static transduction approach, with distinct advantages, such as superior efficiency, reduced timelines required for handling large numbers of cells simultaneously, and faster cell recovery in both target cell lines. This approach curtailed experimental complexity and facilitated a streamlined workflow for a genome-wide screen.

Pooled CRISPR screens typically aim for a balanced transduction efficiency, ranging from 20% to 50%, which corresponds to a multiplicity of infection (MOI) of 0.3 – 0.5. This range ensures that a substantial proportion of cells are transduced with at least one virion, minimizing the number of uninfected cells, while reducing the likelihood of multiple lentiviral integration events^36,82^. However, as our ddPCR analysis highlighted, at transduction efficiencies nearing 40%, the average copy number approaches two copies per cell.

Therefore, in lentiviral delivery-based applications, such as pooled CRISPR screens, where control of copy number is crucial, neglecting cell line-specific responses to transduction, as well as the impact of the experimental setup, can lead to higher-than-expected copy number variation. Such variation can confound data interpretation by introducing unintended multiple perturbations or failing to achieve the intended delivery outcome^38^. Our data strongly support the necessity of determining the required MOI for each target cell model and experimental setup, allowing for a tailored approach that balances transduction efficiency and integration copy number.

In conclusion, our study addresses critical challenges in methodological protocols and scalability, offering valuable insights for designing effective genome-wide CRISPR library delivery using lentiviral vectors in hard-to-transduce suspension cell lines.

## Acknowledgments

We sincerely thank the Borth group at BOKU University for their support and camaraderie throughout this work. We thank Ursula Kiesswetter and Zerina Arnautalic for their contributions to lab coordination and organization. We also extend our gratitude to the Wozniak-Knopp and Grillari groups for providing the HEK293 cell lines used in this study. We thank the BOKU Core Facilities Biomolecular & Cellular Analysis (BmCA) and Food and Bioprocessing at BOKU University, Vienna, Austria. This work was supported by the Austrian Center of Industrial Biotechnology (acib), a COMET center supported by the Austrian Research Promotion Agency FFG and funded by BMVIT, BMDW, SFG, Standortagentur Tirol, Government of Lower Austria, and Vienna Business Agency.

